# plyranges: A grammar of genomic data transformation

**DOI:** 10.1101/327841

**Authors:** Stuart Lee, Dianne Cook, Michael Lawrence

## Abstract

The Bioconductor project provides many interoperable data abstractions for analyzing high-throughput genomics experiments; however implementing a typical genomic workflow with Bioconductor requires learning these abstractions and understanding them at an integrative level. This places a large cognitive burden on the user, especially for non-programmers. To reduce this burden we have created a grammar of genomic data transformation that operates on a single, central Bioconductor data structure, GRanges, which naturally represents genomic intervals and their associated measurements. The grammar defines verbs for performing actions on and between genomic interval data through a simplified, coherent interface to existing Bioconductor infrastructure, resulting in fluent analysis workflows. We have implemented this grammar as an R/Bioconductor package called `plyranges`.

## Background

High-throughput genomics promises to unlock new disease therapies, and strengthen our knowledge of basic biology. To deliver on those promises, scientists must derive a stream of knowledge from a deluge of data. Genomic data is challenging in both scale and complexity. Innovations in sequencing technology often outstrip our capacity to process the output. Beyond their common association with genomic coordinates, genomic data are heterogeneous, consisting of raw sequence read alignments, genomic feature annotations like genes and exons, and summaries like coverage vectors, ChIP-seq peak calls, variant calls, and per-feature read counts. Genomic scientists need software tools to wrangle the different types of data, process the data at scale, test hypotheses, and generate new ones, all while focusing on the biology, not the computation. For the tool developer, the challenge is to define ways to model and operate on the data that align with the mental model of scientists, and to provide an implementation that scales with their ambition.

Several domain specific languages (DSLs) enable scientists to process and reason about heterogeneous genomics data by expressing common operations, such as range manipulation and overlap-based joins, using the vocabulary of genomics. Their implementations either delegate computations to a database, or operate over collections of files in standard formats like BED. An example of the former is the Genome Query Language (GQL) and its distributed implementation GenAp which use a SQL-like syntax for fast retrieval of information of unprocessed sequencing data [1, 2]. Similarly, the Genometric Query Language (GMQL) implements a DSL for combining genomic datasets [3]. The command line application BEDtools develops an extensive algebra for performing arithmetic between two or more sets of genomic regions [4]. All of the aforementioned DSLs are designed to be evaluated either at the command line or embedded in scripts for batch processing. They exist in a sparse ecosystem, mostly consisting of UNIX and database tools that lack biological semantics and operate at the level of files and database tables.

The Bioconductor/R packages `IRanges` and `GenomicRanges` [5, 6, 7] define a DSL for analyzing genomics data with R, an interactive data analysis environment that encourages reproducibility and provides high-level abstractions for manipulating, modelling and plotting data, through state of the art methods in statistical computing. The packages define object-oriented (OO) abstractions for representing genomic data and enable interoperability by allowing users and developers to use these abstractions in their own code and packages. Other genomic DSLs that are embedded in programming languages include pybedtools and valr [8, 9], however these packages lack the interoperability provided by the aforementioned Bioconductor packages and are not easily extended.

The Bioconductor infrastructure models the genomic data and operations from the perspective of the power user, one who understands and wants to take advantage of the subtle differences in data types. This design has enabled the development of sophisticated tools, as evidenced by the hundreds of packages depending on the framework. Unfortunately, the myriad of data structures have overlapping purposes and important but obscure differences in behavior that often confuse the typical end user.

Recently, there has been a concerted, community effort to standardize R data structures and workflows around the notion of tidy data [10]. A tidy dataset is defined as a tabular data structure that has observations as rows and columns as variables, and all measurements pertain to a single observational unit. The tidy data pattern is useful because it allows us to see how the data relate to the design of an experiment and the variables measured. The `dplyr` package [11] defines an application programming interface (API) that maps notions from the general relational algebra to verbs that act on tidy data. These verbs can be composed together on one or more tidy datasets with the pipe operator from the `magrittr` package [12]. Taken together these features enable a user to write human readable analysis workflows.

We have created a genomic DSL called `plyranges` that reformulates notions from existing genomic algebras and embeds them in R as a genomic extension of `dplyr`. By analogy, `plyranges` is to the genomic algebra, as `dplyr` is to the relational algebra. The `plyranges` Bioconductor package implements the language on top of a key subset of Bioconductor data structures and thus fully integrates with the Bioconductor framework, gaining access to its scalable data representations and sophisticated statistical methods.

## Results

### Genomic Relational Algebra

#### Data Model

**Figure 1:**
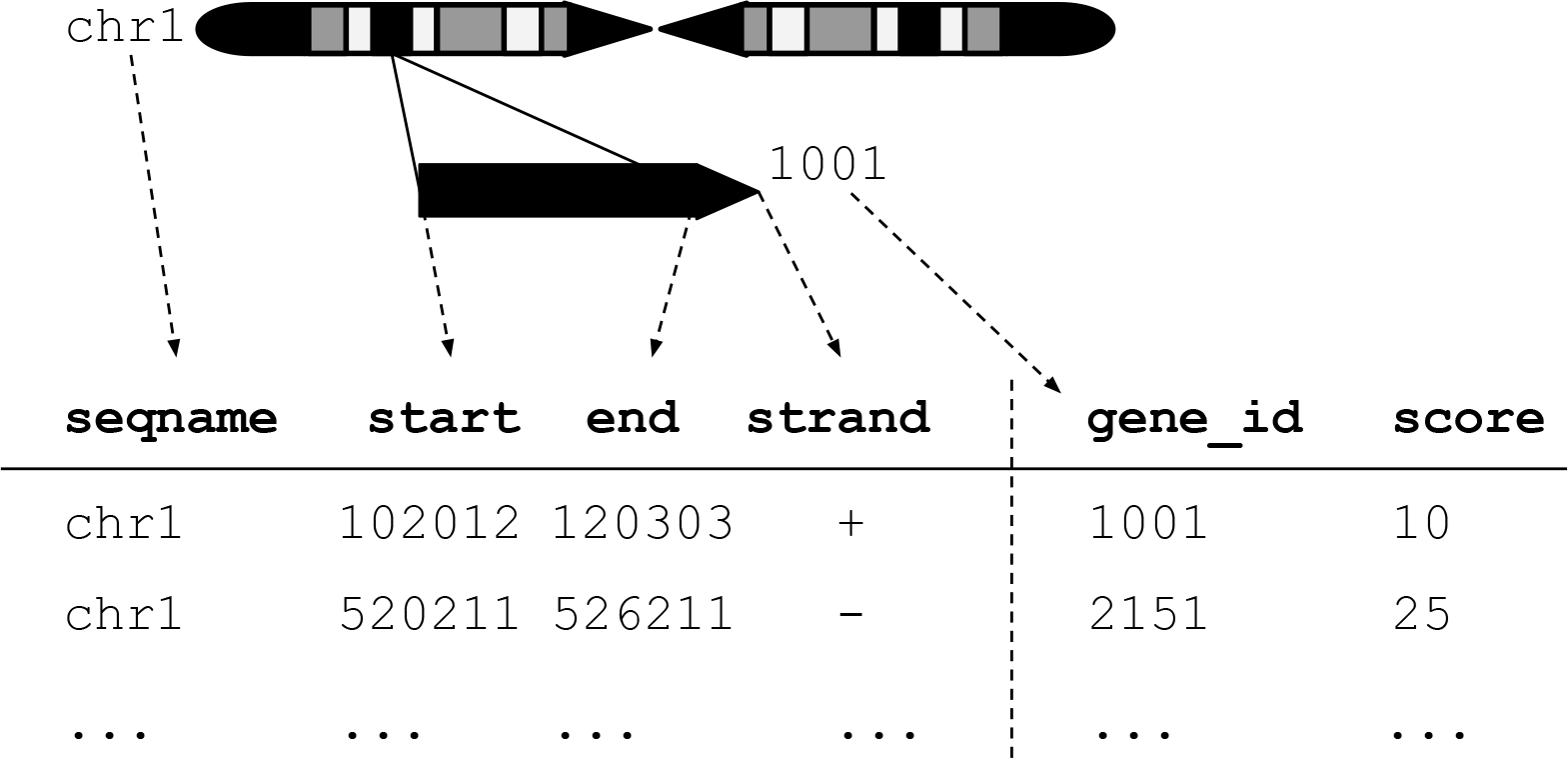
An illustration of the GRanges data model for a sample from an RNA-seq experiment. The core components of the data model include a seqname column (representing the chromosome), a ranges column which consists of start and end coordinates for a genomic region, and a strand identifier (either positive, negative, or un-stranded). Metadata are included as columns to the right of the dotted line as annotations (gene_id) or range level covariates (score).

The `plyranges` DSL is built on the core Bioconductor data structure GRanges, which is a constrained table, with fixed columns for the chromosome, start and end coordinates, and the strand, along with an arbitrary set of additional columns, consisting of measurements or metadata specific to the data type or experiment (figure 1). GRanges balances flexibility with formal constraints, so that it is applicable to virtually any genomic workflow, while also being semantically rich enough to support high-level operations on genomic ranges. As a core data structure, GRanges enables interoperability between `plyranges` and the rest of Bioconductor. Adhering to a single data structure simplifies the API and makes it easier to learn and understand, in part because operations become endomorphic, i.e., they return the same type as their input.

GRanges follow the intuitive tidy data pattern: it is a rectangular table corresponding to a single biological context. Each row contains a single observation and each column is a variable describing the observations. GRanges specializes the tidy pattern in that the observations always pertain to some genomic feature, but it largely remains compatible with the general relational operations defined by `dplyr`. Thus, we define our algebra as an extension of the `dplyr` algebra, and borrow its syntax conventions and design principles.

#### Algebraic operations

The `plyranges` DSL defines an expressive algebra for performing genomic operations with and between GRanges objects (see table 1). The grammar includes several classes of operation that cover most use cases in genomics data analysis. There are range arithmetic operators, such as for resizing ranges or finding their intersection, and operators for merging, filtering and aggregating by range-specific notions like overlap and proximity.

**Table 1:**
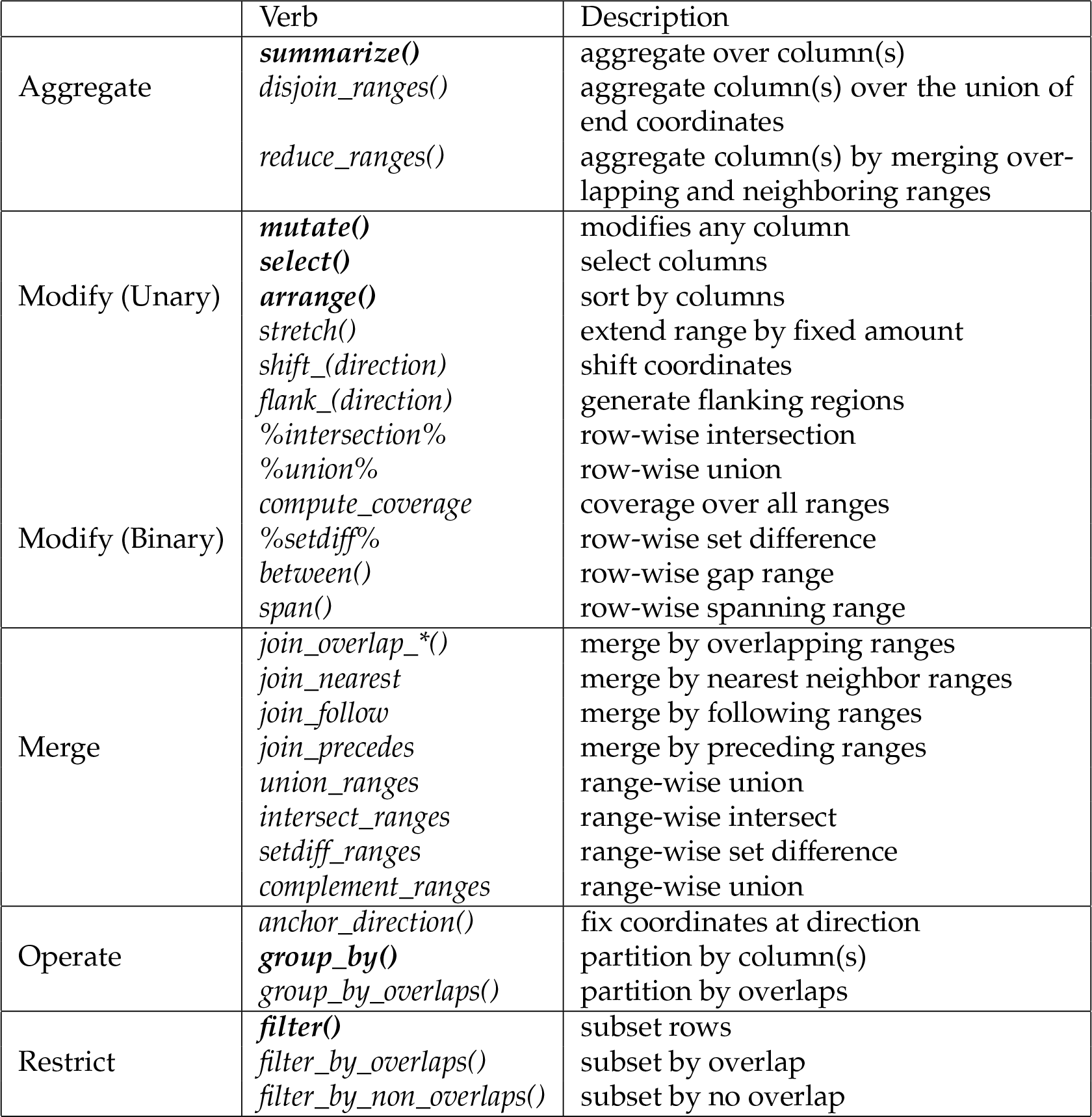
Overview of the plyranges grammar. The core verbs are briefly described and categorized into one of the following higher level categories: aggregate, modify, merge, operate, or restrict. A verb is given bold text if its origin is from the dplyr grammar.

Arithmetic operations transform range coordinates, as defined by their *start*, *end* and *width*. The three dimensions are mutually dependent and partially redundant, so direct manipulation of them is problematic. For example, changing the *width* column needs to change either the *start*, *end* or both to preserve integrity of the object. We introduce the *anchor* modifier to disambiguate these adjustments. Supported anchor points include the start, end and midpoint, as well as the 3’ and 5’ ends for strand-directed ranges. For example, if we anchor the start, then setting the width will adjust the end while leaving the start stationary.

The algebra also defines conveniences for relative coordinate adjustments: *shift* (unanchored adjustment to both start and end) and *stretch* (anchored adjustment of width). We can perform any relative adjustment by some combination of those two operations. The *stretch* operation requires an anchor and assumes the midpoint by default. Since *shift* is unanchored, the user specifies a suffix for indicating the direction: left/right or, for stranded features, upstream/downstream. For example, *shift_right* shifts a range to the right.

The *flank* operation generates new ranges that are adjacent to existing ones. This is useful, for example, when generating upstream promoter regions for genes. Analogous to *shift*, a suffix indicates the side of the input range to flank.

As with other genomic grammars, we define set operations that treat ranges as sets of integers, including *intersect*, *union*, *difference*, and *complement*. There are two sets of these: parallel and merging. For example, the parallel intersection (*x %intersect% y*) finds the intersecting range between *xi* and *yi* for *i* in *1… n*, where *n* is the length of both *x* and *y*. In contrast, the merging intersection (*intersect_ranges(x, y)*) returns a new set of disjoint ranges representing wherever there was overlap between a range in *x* and a range in *y*. Finding the parallel union will fail when two ranges have a gap, so we introduce a *span* operator that takes the union while filling any gap. The *complement* operation is unique in that it is unary. It finds the regions not covered by any of the ranges in a single set. Closely related is the *between* parallel operation, which finds the gap separating *xi* and *yi*. The binary operations are callable from within arithmetic, restriction and aggregation expressions.

To support merging, our algebra recasts finding overlaps or nearest neighbors between two genomic regions as variants of the relational join operator. A join acts on two GRanges objects: *x* and *y*. The join operator is relational in the sense that metadata from the *x* and *y* ranges are retained in the joined range. All join operators in the `plyranges` DSL generate a set of hits based on overlap or proximity of ranges and use those hits to merge the two datasets in different ways. There are four supported matching algorithms: *overlap*, *nearest*, *precede*, and *follow* (figure 2). We can further restrict the matching by whether the query is completely *within* the subject, and adding the *directed* suffix ensures that matching ranges have the same direction (strand).

**Figure 2:**
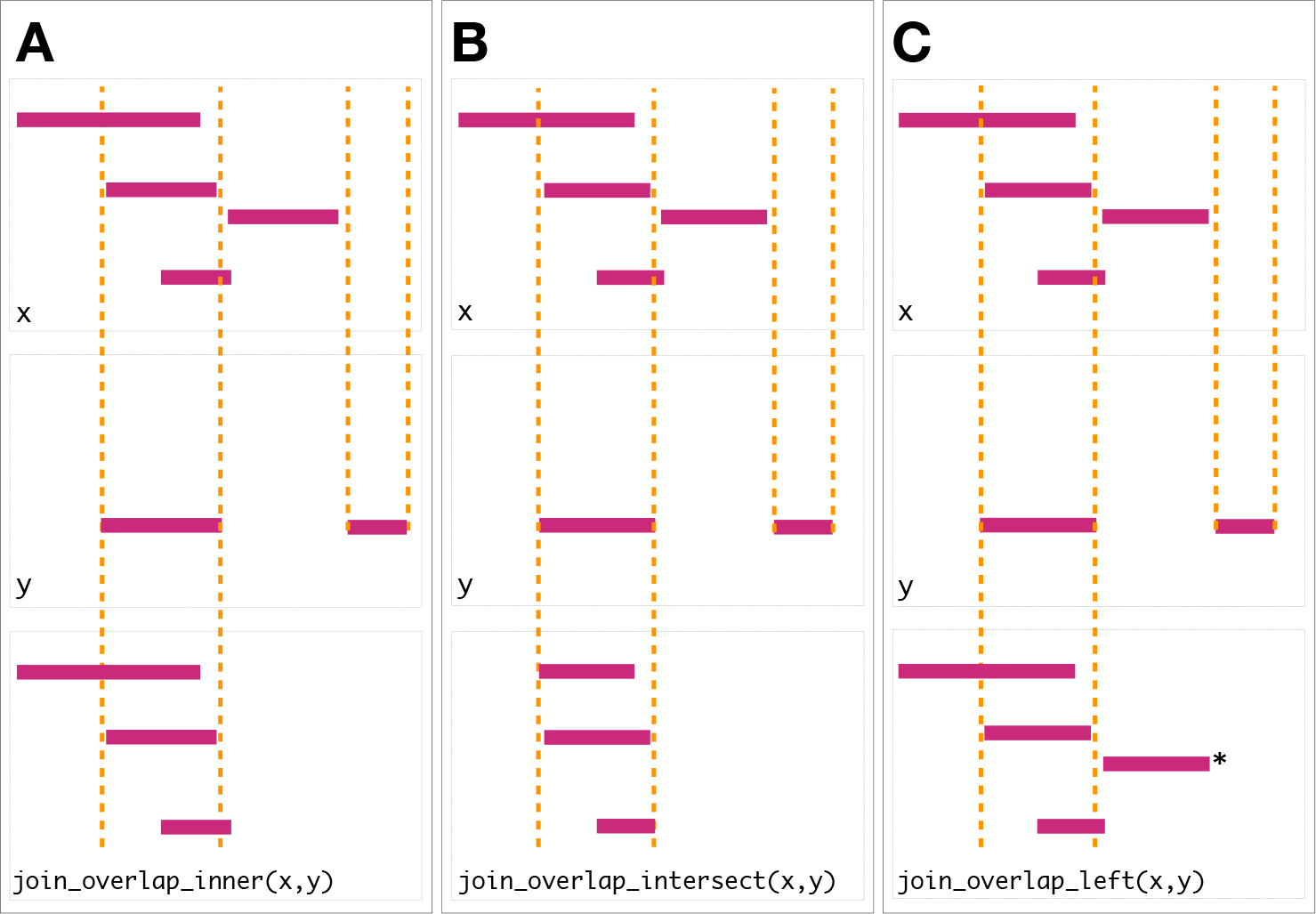
Illustration of the three overlap join operators. Each join takes two GRanges objects, *x* and *y* as input. A ‘Hits’ object for the join is computed which consists of two components. The first component contains the indices of the ranges in *x* that have been overlapped (the rectangles of *x* that cross the orange lines). The second component consists of the indices of the ranges in *y* that overlap the ranges in *x*. In this case a range in *y* overlaps the ranges in *x* three times, so the index is repeated three times. The resulting ‘Hits’ object is used to modify *x* by where it was ‘hit’ by *y* and merge all metadata columns from *x* and *y* based on the indices contained in the ‘Hits’ object. This procedure is applied generally in the `plyranges` DSL for both overlap and nearest neighbor operations. The join semantics alter what is returned: **A**: for an **inner** join the *x* ranges that are overlapped by *y* are returned. The returned ranges also include the metadata from the *y* range that overlapped the three *x* ranges. **B** An **intersect** join is identical to an inner join except that the intersection is taken between the overlapped *x* ranges and the *y* ranges. **C** For the **left** join all *x* ranges are returned regardless of whether they are overlapped by *y*. In this case the third range (rectangle with the asterisk next to it) of the join would have missing values on metadata columns that came from *y*.

For merging based on the hits, we have three modes: *inner*, *intersect* and *left*. The *inner* overlap join is similar to the conventional inner join in that there is a row in the result for every match. A major difference is that the matching is not by identity, so we have to choose one of the ranges from each pair. We always choose the left range. The *intersect* join uses the intersection instead of the left range. Finally, the overlap *left* join is akin to left outer join in Codd’s relational algebra: it performs an overlap inner join but also returns all *x* ranges that are not hit by the *y* ranges.

**Figure 3:**
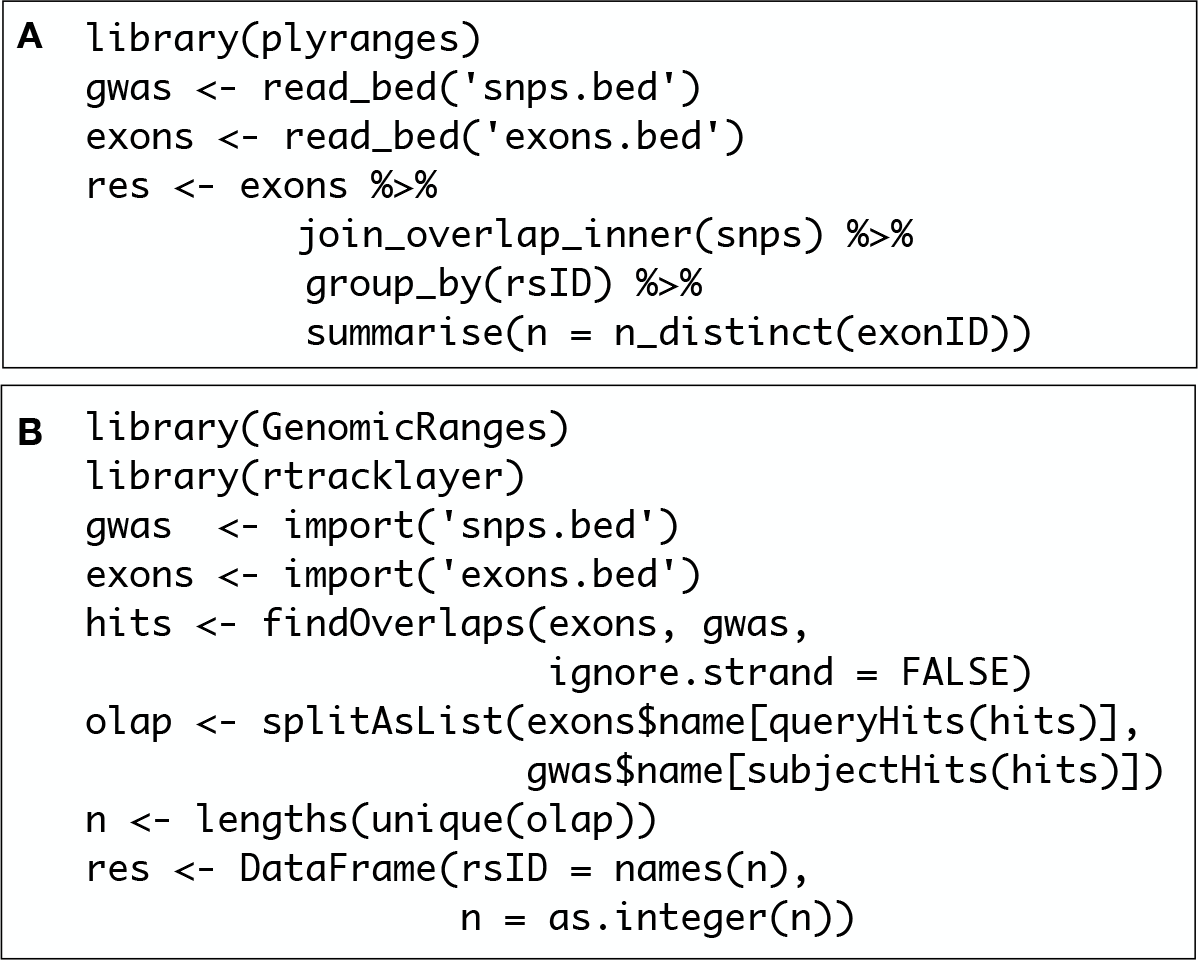
Idiomatic code examples for `plyranges` (**A**) and `GenomicRanges` (**B**) illustrating an overlap and aggregate operation that returns the same result. In each example, we have two BED files consisting of SNPs that are genome-wide association study (GWAS) hits and reference exons. Each code block counts for each SNP the number of distinct exons it overlaps. The `plyranges` code achieves this with an overlap join followed by partitioning and aggregation. Strand is ignored by default here. The `GenomicRanges` code achieves this using the ‘Hits’ and ‘List’ classes and their methods.

Since the GRanges object is a tabular data structure, our grammar includes operators to filter, sort and aggregate by columns in a GRanges. These operations can be performed over partitions formed using the *group_by* modifier. Together with our algebra for arithmetic and merging, these operations conform to the semantics and syntax of the dplyr grammar. Consequently, `plyranges` code is generally more compact than the equivalent `GenomicRanges` code (figure 3).

### Developing workflows with plyranges

Here we provide illustrative examples of using the `plyranges` DSL to show how our grammar could be integrated into genomic data workflows. As we construct the work-flows we show the data output intermittently to assist the reader in understanding the pipeline steps. The workflows highlight how interoperability with existing Bioconductor infrastructure, enables easy access to public datasets and methods for analysis and visualization.

### Peak Finding

In the workflow of ChIP-seq data analysis, we are interested in finding peaks from islands of coverage over chromosome. Here we will use `plyranges` to call peaks from islands of coverage above 8 then plot the region surrounding the tallest peak.

Using `plyranges` and the the Bioconductor package `AnnotationHub` [13] we can download and read BigWig files from ChIP-Seq experiments from the Human Epigenome Roadmap project [14]. Here we analyse a BigWig file corresponding to H3 lysine 27 trimethylation (H3K27Me3) of primary T CD8+ memory cells from peripheral blood, focussing on coverage islands over chromosome 10.

First, we extract the genome information from the BigWig file and filter to get the range for chromosome 10. This range will be used as a filter when reading the file.

`library(plyranges)`

`chr10_ranges <- bw_file %>%`

`get_genome_info() %>%`

`filter(seqnames == "chr10")`

Then we read the BigWig file only extracting scores if they overlap chromosome 10. We also add the genome build information to the resulting ranges. This book-keeping is good practice as it ensures the integrity of any downstream operations such as finding overlaps.

`chr10_scores <- bw_file %>%`

`read_bigwig(overlap_ranges = chr10_ranges) %>%`

`set_genome_info(genome = "hg19")`

`chr10_scores`

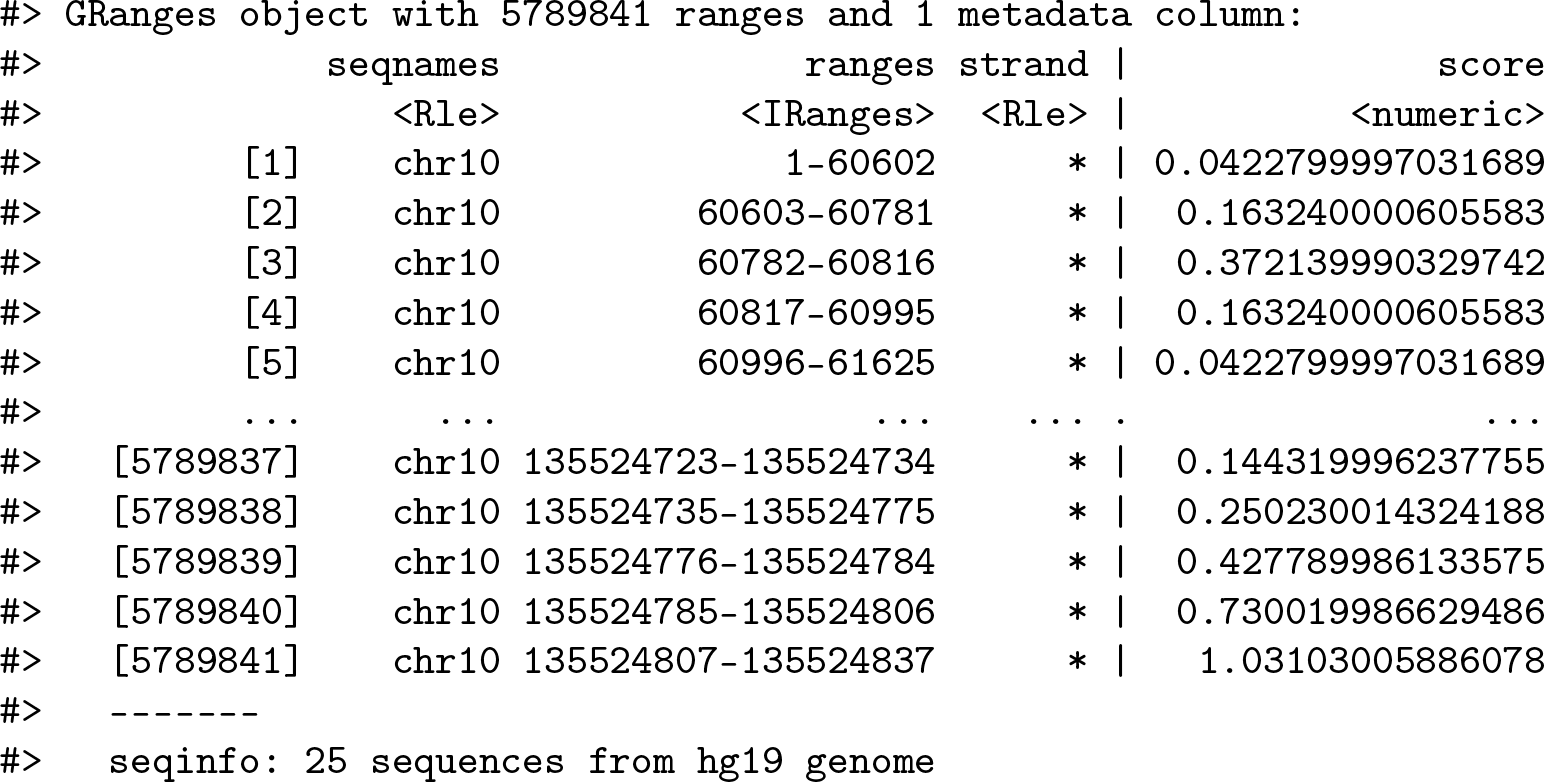

We then filter for regions with a coverage score greater than 8, and following this reduce individual runs to ranges representing the islands of coverage. This is achieved with the `reduce_ranges`() function, which allows a summary to be computed over each island: in this case we take the maximum of the scores to find the coverage peaks over chromosome 10.

`all_peaks <- chr10_scores %>%`

`filter(score > 8) %>%`

`reduce_ranges(score = max(score))`

`all_peaks`

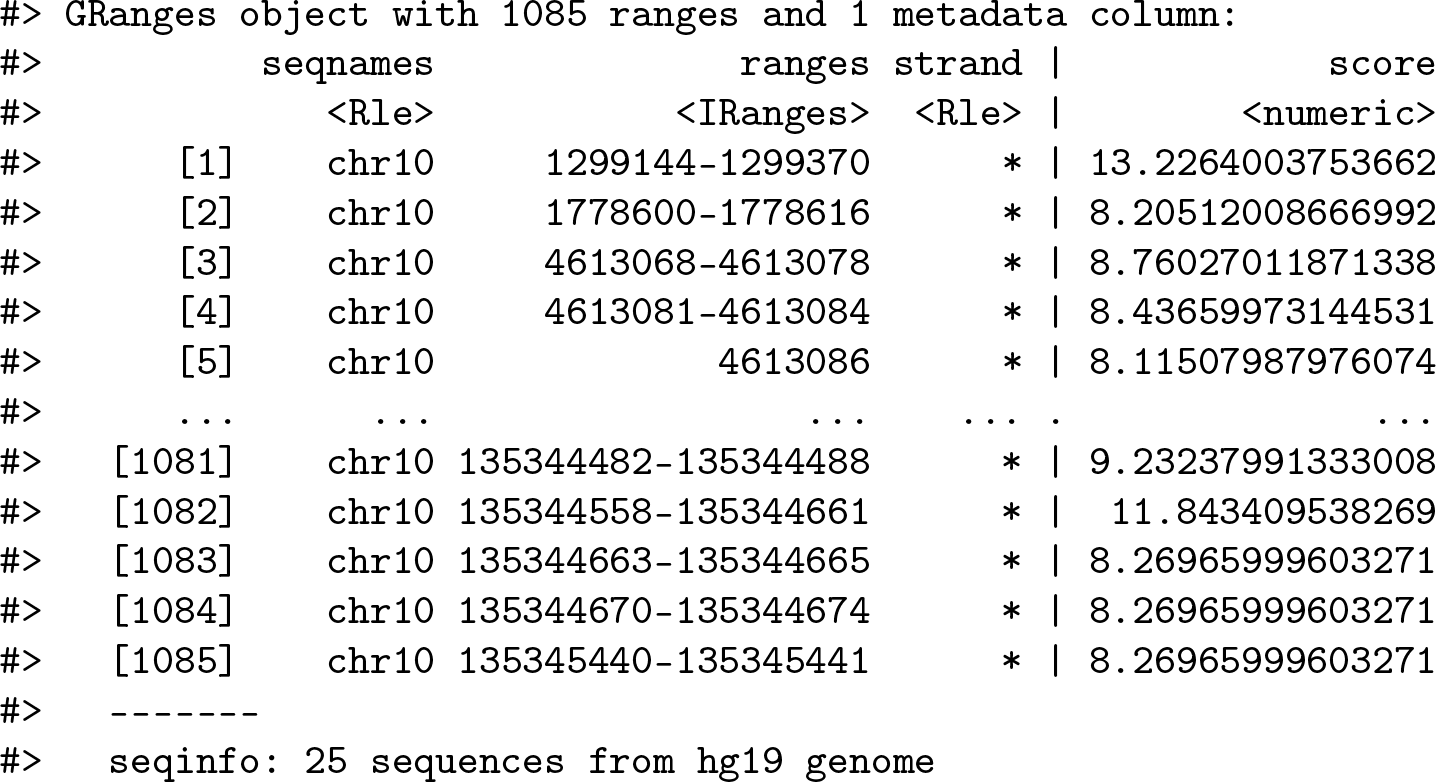

Returning to the GRanges object containing normalized coverage scores, we filter to find the coordinates of the peak containing the maximum coverage score. We can then find a 5000 nt region centered around the maximum position by anchoring and modifying the width.

`chr10_max_score_region <- chr10_scores %>%`

`filter(score == max(score)) %>%`

`anchor_center() %>%`

`mutate(width = 5000)`

Finally, the overlap inner join is used to restrict the chromosome 10 coverage islands, to the islands that are contained in the 5000nt region that surrounds the max peak (figure 4).

`peak_region <- chr10_scores %>%`

`join_overlap_inner(chr10_max_score_region)`

`peak_region`

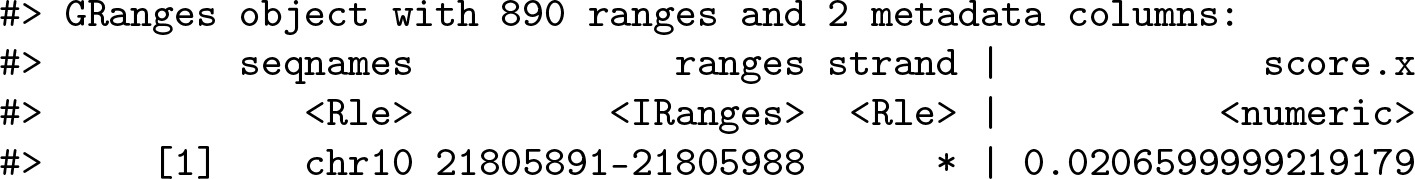

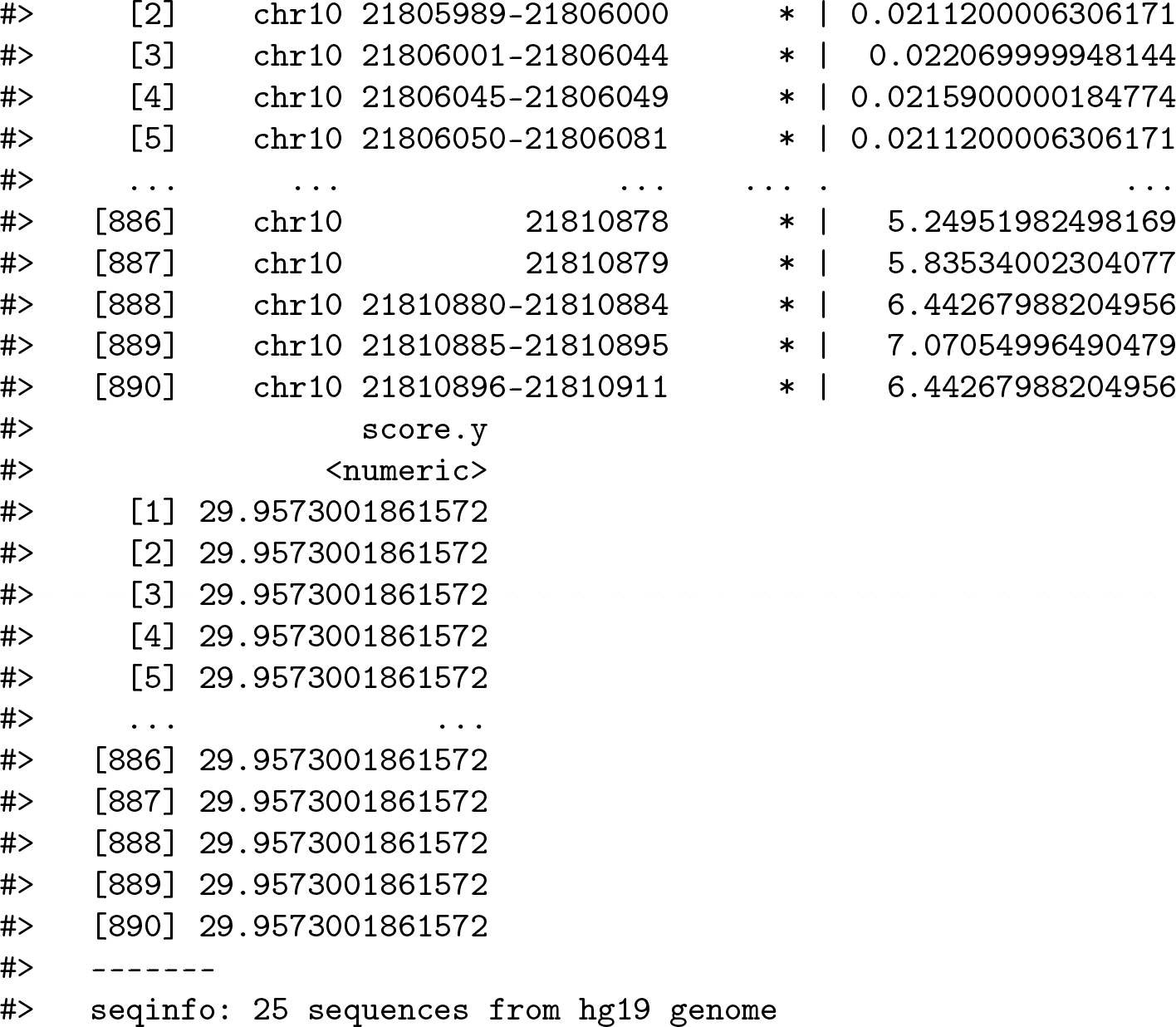

**Figure 4:**
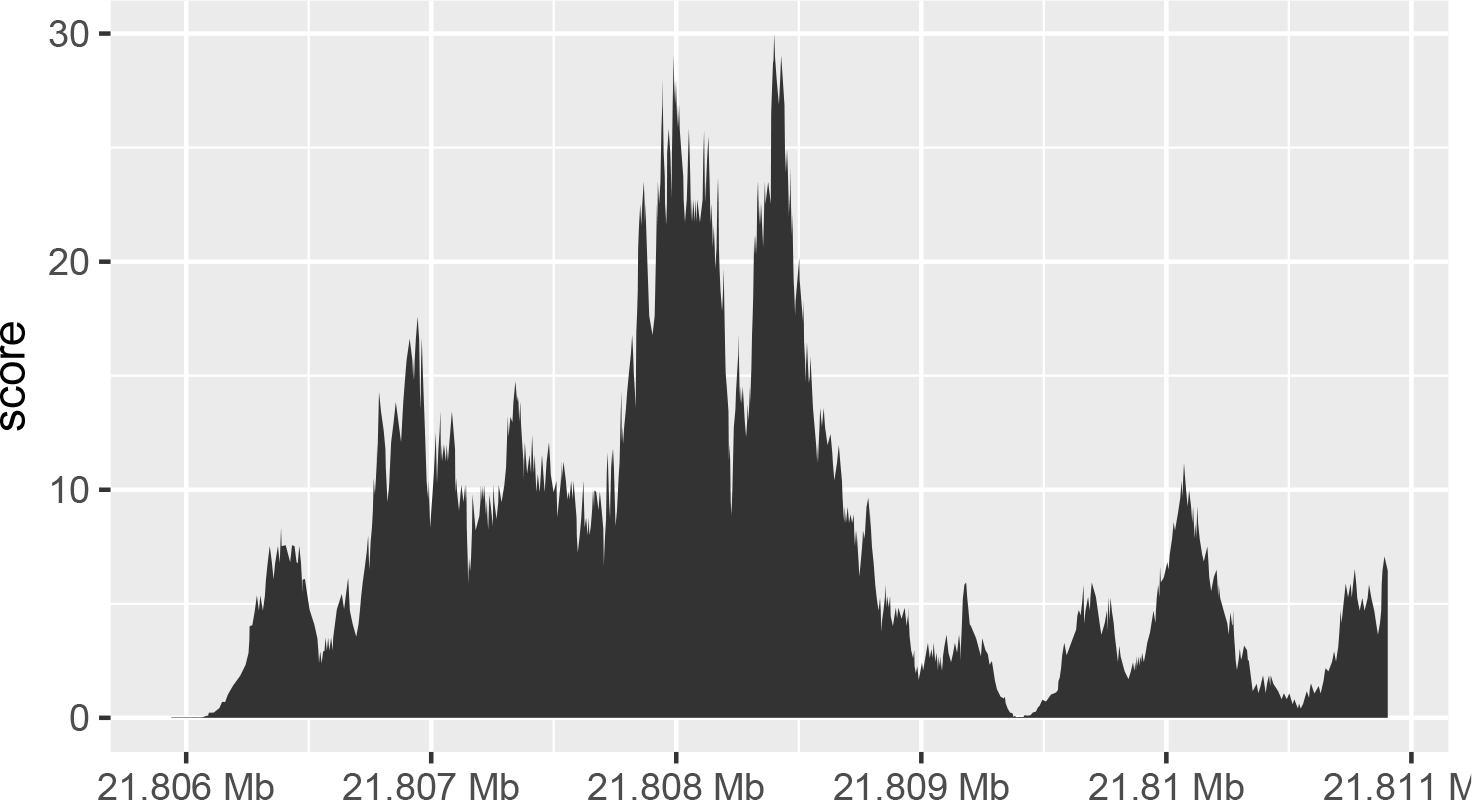
The final result of the `plyranges` operations to find a 5000nt region surrounding the peak of normalised coverage scores over chromosome 10, displayed as a density plot.

### Computing Windowed Statistics

Another common operation in genomics data analysis is to compute data summaries over genomic windows. In `plyranges` this can be achieved via the `group_by_overlaps`() operator. We bin and count and find the average GC content of reads from a H3K27Me3 ChIP-seq experiment by the Human Epigenome Roadmap Consortium.

We can directly obtain the genome information from the header of the BAM file: in this case the reads were aligned to the hg19 genome build and there are no reads overlapping the mitochondrial genome.

`locations <- h1_bam_sorted %>%`

`read_bam() %>%`

`get_genome_info()`

Next we only read in alignments that overlap the genomic locations we are interested in and select the query sequence. Note that the reading of the BAM file is deferred: only alignments that pass the filter are loaded into memory. We can add another column representing the GC proportion for each alignment using the `letterFrequency`() function from the `Biostrings` package [15]. After computing the GC proportion as the score column, we drop all other columns in the GRanges object.

`alignments <- h1_bam_sorted %>%`

`read_bam() %>%`

`filter_by_overlaps(locations) %>%`

`select(seq) %>%`

`mutate(`

`score = as.numeric(letterFrequency(seq, "GC", as.prob = TRUE))`

`) %>%`

`select(score)`

`alignments`

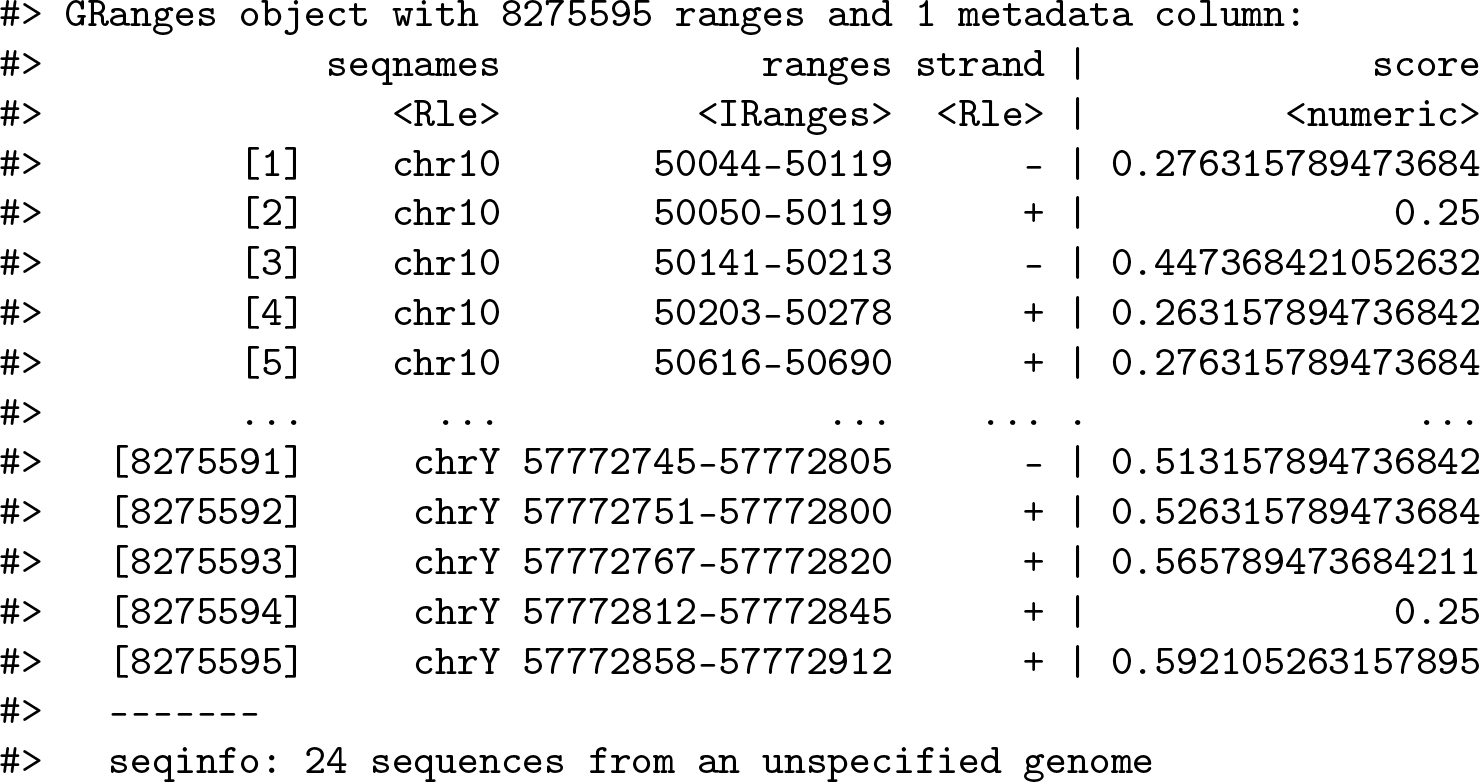

Finally, we create 10000nt tiles over the genome and compute the number of reads and average GC content over all reads that fall within each tile using an overlap join and merging endpoints.

`bins <- locations %>%`

`tile_ranges(width = 10000L)`

`alignments_summary <- bins %>%`

`join_overlap_inner(alignments) %>%`

`disjoin_ranges(n = n(), avg_gc = mean(score))`

`alignments_summary`

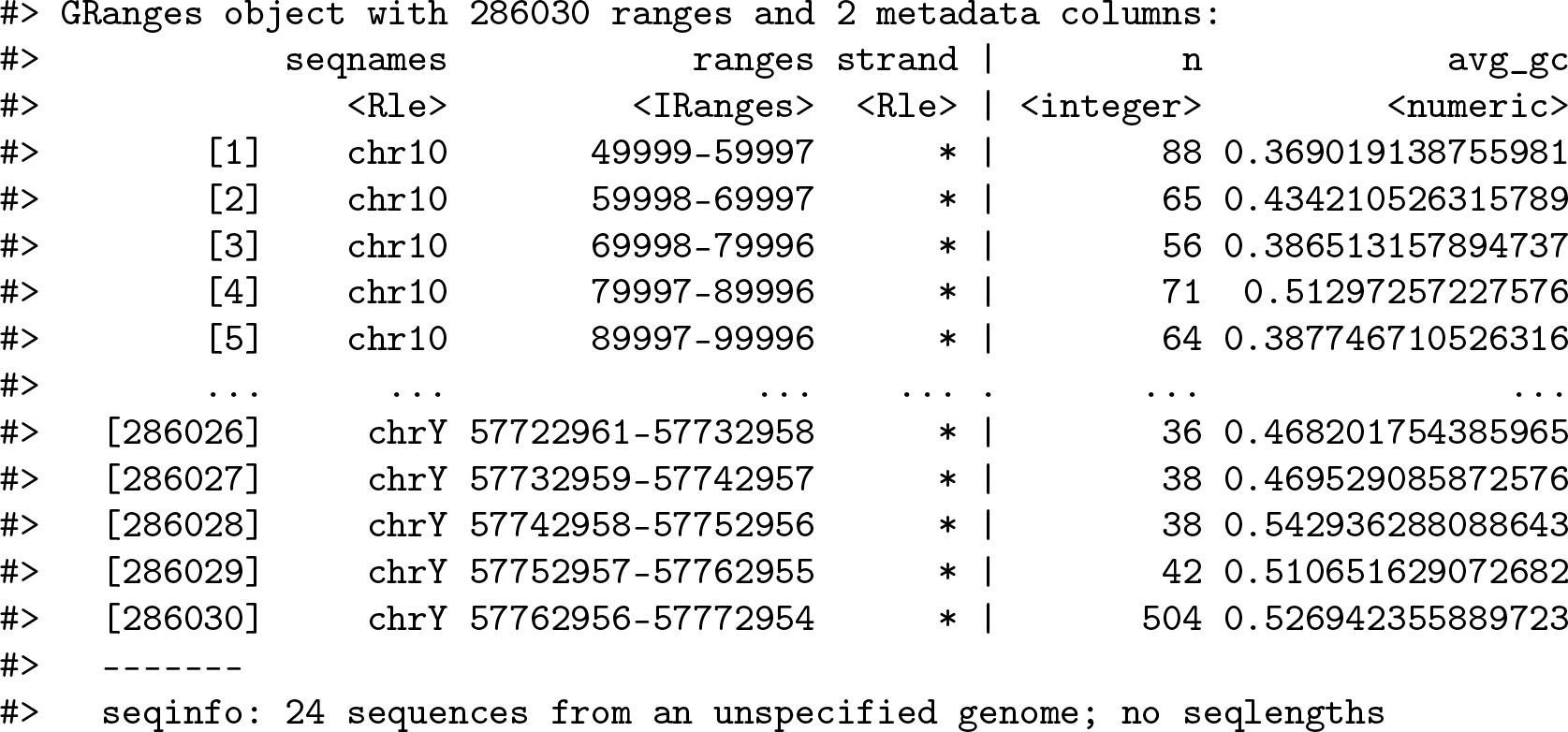

### Quality Control Metrics

We have created a GRanges object from genotyping performed on the H1 cell line, consisting of approximately two million single nucleotide polymorphisms (SNPs) and short insertion/deletions (indels). The GRanges object consists of 7 columns, relating to the alleles of a SNP or indel, the B-allele frequency, log relative intensity of the probes, GC content score over a probe, and the name of the probe. We can use this information to compute the transition-transversion ratio, a quality control metric, within each chromosome in GRanges object.

First we filter out the indels and mitochondrial variants. Then we create a logical vector corresponding to whether there is a transition event.

`h1_snp_array <- h1_snp_array %>%`

`filter(!(ref %in% c("I", "D")), seqnames != "M") %>%`

`mutate(transition = (ref %in% c("A", "G") & alt %in% c("G","A"))|(ref %in% c("C","T") & alt %in% c("T", "C")))`

We then compute the transition-transversion ratio over each chromosome using `group_by`() in combination with `summarize`() (figure 5).

`ti_tv_results <- h1_snp_array %>%`

`group_by(seqnames) %>%`

`summarize(n_snps = n(),ti_tv = sum(transition) / sum(!transition))`

`ti_tv_results`

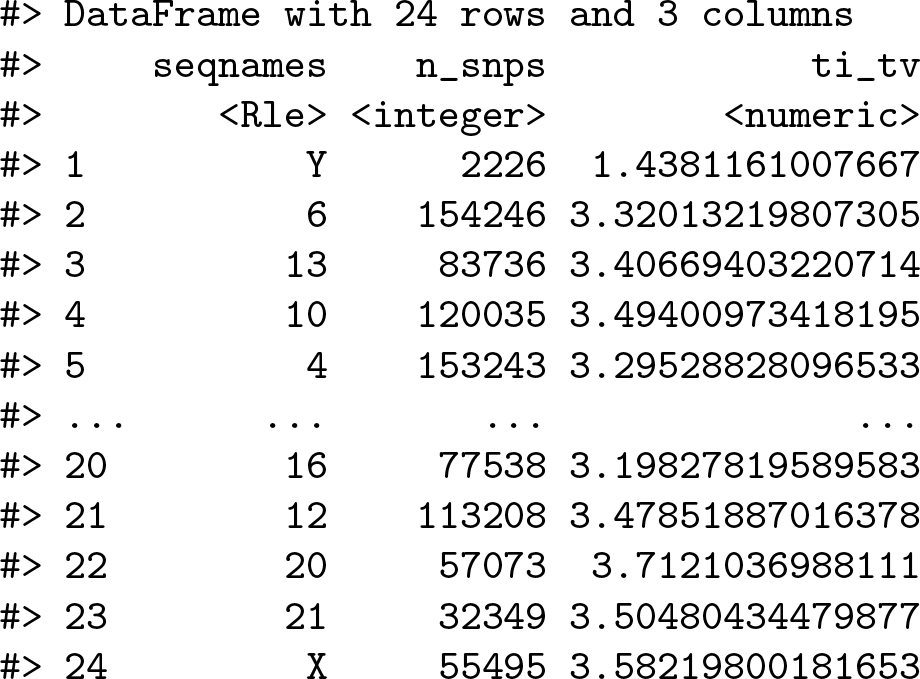

**Figure 5:**
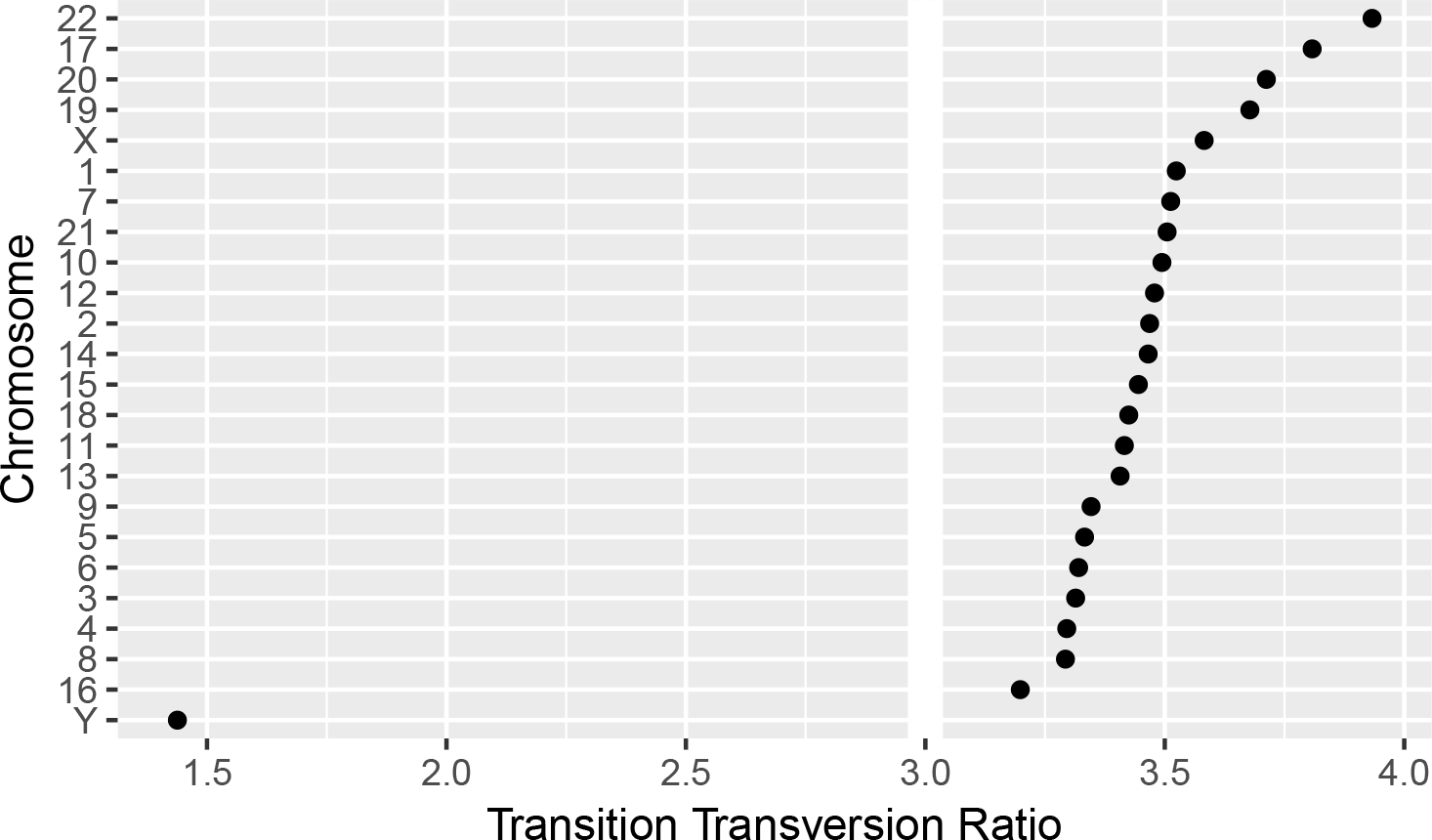
The final result of computing quality control metrics over the SNP array data with `plyranges`, displayed as a dot plot. Chromosomes are ordered by their estimated transition-transversion ratio. A white reference line is drawn at the expected ratio for a human exome.

## Discussion

The design of `plyranges` adheres to well understood principles of language and API design: cognitive consistency, cohesion, expressiveness and endomorphism [16]. To varying degrees, these principles also underlie the design of dplyr and the Bioconductor infrastructure.

We have aimed for `plyranges` to have a simple and direct mapping to the user’s cognitive model, i.e., how the user thinks about the data. This requires careful selection of the level of abstraction so that the user can express workflows in the language of genomics. This motivates the adoption of the tidy GRanges object as our central data structure. The basic data.frame and `dplyr` tibble lack any notion of genomic ranges and so could not easily support our genomic grammar, with its specific verbs for range-oriented data manipulation. Another example of cognitive consistency is how `plyranges` is insensitive to direction/strand by default when, e.g., detecting overlaps. `GenomicRanges` has the opposite behavior. We believe that defaulting to purely spatial overlap is most intuitive to most users.

To further enable cognitive consistency, `plyranges` functions are cohesive. A function is defined to be cohesive if it performs a singular task without producing any side-effects. Singular tasks can always be broken down further at lower levels of abstraction. For example, to resize a range, the user needs to specify which position (start, end, midpoint) should be invariant over the transformation. The `resize`() function from the `GenomicRanges` package has a `fix` argument that sets the anchor, so calling `resize`() coalesces anchoring and width modification. The coupling at the function call level is justified since the effect of setting the width depends on the anchor. However, `plyranges` increases cohesion and decouples the anchoring into its own function call.

Increasing cohesion simplifies the interface to each operation, makes the meaning of arguments more intuitive, and relies on function names as the primary means of expression, instead of a more complex mixture of function and argument names. This results in the user being able to conceptualize the `plyranges` DSL as a flat catalog of functions, without having to descend further into documentation to understand a function’s arguments. A flat function catalog also enhances API discoverability, particularly through auto-completion in integrated developer environments (IDEs). One downside of pushing cohesion to this extreme is that function calls become coupled, and care is necessary to treat them as a group when modifying code.

Like `dplyr, plyranges` verbs are functional: they are free of side effects and are generally endomorphic, meaning that when the input is a GRanges object they return a GRanges object. This enables chaining of verbs through syntax like the forward pipe operator from the `magrittr` package. This syntax has a direct cognitive mapping to natural language and the intuitive notion of pipelines. The low-level object-oriented APIs of Bioconductor tend to manipulate data via sub-replacement functions, like `start(gr) <- x`. These ultimately produce the side effect of replacing a symbol mapping in the current environment and thus are not amenable to so-called fluent syntax.

Expressiveness relates to the information content in code: the programmer should be able to clarify intent without unnecessary verbosity. For example, our overlap-based join operations are more concise than the multiple steps necessary to achieve the same effect in the original `GenomicRanges` API. In other cases, the `plyranges` API increases verbosity for the sake of clarity and cohesion. Explicitly calling *anchor()* can require more typing, but the code is easier to comprehend. Another example is the set of routines for importing genomic annotations, including *read_gff()*, *read_bed()*, and *read_bam()*. Compared to the generic *import()* in `rtracklayer`, the explicit format-based naming in `plyranges` clarifies intent and the type of data being returned. Similarly, every `plyranges` function that computes with strand information indicates its intentions by including suffixes such as *directed*, *upstream* or *downstream* in its name, otherwise strand is ignored. The `GenomicRanges` API does not make this distinction explicit in its function naming, instead relying on a parameter that defaults to strand sensitivity, an arguably confusing behavior.

The implementation of `plyranges` is built on top of Bioconductor infrastructure, meaning most functions are constructed by composing generic functions from core Bioconductor packages. As a result, any Bioconductor packages that uses data structures that inherit from GRanges will be able to use `plyranges` for free. Another consequence of building on top of Bioconductor generics is that the speed and memory usage of `plyranges` functions are similar to the highly optimized methods implemented in Bio-conductor for GRanges objects.

A caveat to constructing a compatible interface with `dplyr` is that `plyranges` makes extensive use of non-standard evaluation in R via the `rlang` package [17]. Simply, this means that computations are evaluated in the context of the GRanges objects. Both `dplyr` and `plyranges` are based on the `rlang` language, because it allows for more expressive code that is free of repeated references to the container. Implicitly referencing the container is particularly convenient when programming interactively. Consequently, when programming with `plyranges`, a user needs to generally understand the `rlang` language and how to adapt their code accordingly. Users familiar with the tidyverse should already have such knowledge.

## Conclusion

We have shown how to create expressive and reproducible genomic workflows using the `plyranges` DSL. By realising that the GRanges data model is tidy we have highlighted how to implement a grammar for performing genomic arithmetic, aggregation, restriction and merging. Our examples show that `plyranges` code is succinct, human readable and can take advantage of the interoperability provided by the Bioconductor ecosystem and the R language.

We also note that the grammar elements and design principles we have described are programming language agnostic and could be easily be implemented in another language where genomic information could be represented as a tabular data structure. We chose R because it is what we are familiar with and because the aforementioned Bioconductor packages have implemented the GRanges data structure.

We aim to continue developing the `plyranges` package and to extend it for use with more complex data structures, such as the SummarizedExperiment class, the core Bioconductor data structure for representing experimental results (e.g., counts) from multiple sample experiments in conjunction with feature and sample metadata. Although, the SummarizedExperiment is not strictly tidy, it does consist of three tidy data structures that are related by feature and sample identifiers. Therefore, the grammar and design of the `plryanges` DSL is naturally extensible to the SummarizedExperiment.

As the `plyranges` interface encourages tidy data practices, it integrates well with the grammar of graphics [18]. To achieve responsive performance, interactive graphics rely on lazy data access and computing patterns, so the deferred mechanisms within `lyranges` should help support interactive genomics applications.

The `plyranges` package can be obtained via the Bioconductor project website https://bioconductor.org or accessed via Github https://github.com/sa-lee/plyranges.

## Methods

### Data Availability

The BigWig file for the H3K27Me3 primary T CD8+ memory cells from peripheral blood ChIP-seq data was downloaded from the AnnotationHub package (2.12.0) under accession AH33458. The BAM file corresponding to the H1 cell line ChIP-seq data is available at GEO under accession GSM433167. The SNP array data for the H1 cell line data is available at GEO under accession GSM1463263.

### Software Versions

To produce the workflows as described in results section we used R version 3.5 with the development version of plyranges (1.1.10) and the BioStrings package (2.48.0) installed.

All code required to reproduce this article is available at https://github.com/sa-lee/plyranges-paper.

## Acknowledgements

We would like to thank Dr Matthew Ritchie at the Walter and Eliza Hall Institute and Dr Paul Harrison for their feedback on earlier drafts of this work. We would also like to thank Lori Shepherd and Hèrve Pages for the code review they performed. This article was written with knitr [19] and the figures were made with `ggbio` [20].

